# Hippocampal silent infarct leads to subtle cognitive decline that is associated with inflammation and gliosis at twenty-four hours after injury in a rat model

**DOI:** 10.1101/2020.09.29.318204

**Authors:** Caitlin A. Finney, Margaret J. Morris, R. Frederick Westbrook, Nicole M. Jones

## Abstract

Silent infarcts (SI) are subcortical cerebral infarcts that occur in the absence of clinical symptoms commonly associated with ischemia and are linked to dementia development. Little is known about the pathophysiology underlying the cognitive dysfunction associated with SI, and few studies have examined the early cellular responses and neurobiological underpinnings. We induced SI in adult male Sprague-Dawley rats using an infusion of endothelin-1 in the CA1 dorsal hippocampus. Twenty-four hours later, we assessed cognition using the hippocampal-dependent object place recognition task. We also examined whether the resulting cognitive effects were associated with common markers of ischemia, specifically cell and synapse loss, gliosis, and inflammation, using histology and immunohistochemistry. Hippocampal SI led to subtle cognitive impairment on the object place recognition task 24-hours post-injury. This was characterized by a significant difference in exploration proportion relative to a pre-injury baseline and a positive association between time spent with both the moved and unmoved objects. SI did not result in any detectable cell or synaptophysin loss, but did increase apoptosis, gliosis and inflammation in the CA1. Principal component analysis indicated the main variables associated with hippocampal SI included increased time spent with the unmoved object, gliosis, apoptosis and inflammation as well as decreased exploration proportion and CA1 cells. Our data demonstrate that hippocampal SI can lead to cognitive dysfunction 24-hours after injury. Further, this appears to be driven by early degenerative processes including apoptosis, gliosis and inflammation, suggesting that these may be targets for early interventions treating hippocampal SI and its cognitive consequences.

## 1. Introduction

Silent infarcts (SI) are a type of cerebral small vessel disease that are defined by the presence of cerebral infarction, usually in subcortical areas, in the absence of clinical symptoms commonly associated with ischemia (Bernick et al., 2001; Chopard et al., 2018; Das et al., 2008; Hahne, Monnig, & Samol, 2016). Recent evidence suggests that for every diagnosed clinical stroke at least five SIs occur, with an overall population prevalence rate of 8-28% (Dempsey, Vemuganti, Varghese, & Hermann, 2010; Leary & Saver, 2003; Vermeer et al., 2003a; Vermeer, Longstreth, & Koudstaal, 2007). There are many risk factors for SI, including diabetes, hypertension, atrial fibrillation, previous stroke, and cardiovascular surgery (Akhtar et al., 2018; Braemswig et al., 2013; Chopard et al., 2018; Das et al., 2008; Geerlings et al., 2010; Grimaldi et al., 2017; Hauville, Ben-Dor, Lindsay, Pichard, & Waksman, 2012; Vermeer et al., 2007). Another risk factor is age, with the incidence of SI increasing from 11% in 55-65 year-olds to greater than 50% in those 85 and older (Howard et al., 2007; Vermeer et al., 2003a). Understanding the pathophysiology of SI is especially important in the globally aging population.

There is a growing body of evidence that, despite being traditionally characterized by the absence of overt clinical symptoms, SI leads to a high incidence of cognitive impairment, especially in elderly patients (Hahne et al., 2016; Howard et al., 2007; Saini et al., 2015; Vermeer et al., 2007). Aside from diagnostic imaging, SI may be best diagnosed by cognitive decline in the absence of overt symptoms of disease or disorder (Dempsey et al., 2010; Hernandez et al., 2015). In fact, SI has been proposed as a possible contributor to the strong relationship between cognitive decline and physical health problems including heart failure and atrial fibrillation (Conen et al., 2019; Pressler, 2008). Studies have consistently shown that SI is associated with decline in both overall cognitive function and in specific cognitive domains, including memory, executive function and processing speed (Blum et al., 2012; Capoccia, Sbarigia, Rizzo, Mansour, & Speziale, 2012; Cinier et al., 2018; Debette & Markus, 2010; Makin, Turpin, Dennis, & Wardlaw, 2013; Saini et al., 2012; Schmidt et al., 2004; Squarzoni et al., 2017; Vermeer et al., 2003b; Zhang et al., 2019). The cognitive symptoms of SI may worsen over time (Hahne et al., 2016; Howard et al., 2007; Saini et al., 2015; Vermeer et al., 2007). Indeed, SI is associated with a doubled risk of dementias, including Alzheimer’s disease and vascular dementia, and may predict more severe forms of these diseases (Blum et al., 2012; Chopard et al., 2018; Debette & Markus, 2010; Gianotti et al., 2004; Hauville et al., 2012; Jellinger, 2002; Vermeer et al., 2003b).

The pathophysiological mechanisms underlying the relationship between SI and cognitive decline are poorly understood. Most SI are characterized by lacunar infarcts, which are associated with microstructural abnormalities in the hippocampus and selective deficits in spatial navigation (Masuda, Nabika, & Notsu, 2001; Wu et al., 2016). Importantly, there can be lasting hippocampal tissue damage after SI which is associated with cognitive decline (Faraji, Kurio, & Metz, 2012; Sharma et al., 2019). In elderly patients with SI, hippocampal atrophy is present and associated with cognitive impairment or dementia (Blum et al., 2012; Fein et al., 2000; Jellinger, 2002). Indeed, the presence of SI accounted for 48% of the variance in neuropsychological data obtained from dementia patients, suggesting that SI can lead to hippocampal degeneration and atrophy (Gianotti et al., 2004). Such hippocampal atrophy is thought to emerge from a combination of ischemic events and other degenerative pathologies, as well as an inflammatory response triggered by the microemboli associated with SI (Capoccia et al., 2012; Fein et al., 2000). SI are especially destructive in the CA1 region of the dorsal hippocampus, particularly in pyramidal neurons that have been shown to undergo selective and delayed degeneration (Blum et al., 2012; Calabresi, Centonze, Pisani, Cupini, & Bernardi, 2003).

Much of what is known about the relationship between SI and cognitive decline, as well as the underlying pathophysiology, has come from human population studies. These rely on the use of diagnostic imaging technologies, such as Magnetic Resonance Imaging (MRI), which provide limited insight into pathophysiology apart from indications of gross structural abnormalities (Hahne et al., 2016). They have also used post-mortem tissue which is difficult to source, and can be difficult to interpret in view of pre-mortem confounds such as medication use, impacting our understanding of the neurobiological mechanisms underlying SI and cognitive decline (Lewis, 2002; Stan et al., 2006). These issues highlight the need for clearly defined pre-clinical SI models to facilitate the development and testing of new treatments for SI, which are currently lacking (Sharma et al., 2019; Vermeer et al., 2007). Recent studies have modelled SI in rats through the use of intracerebral infusions of endothelin-1, a potent vasoconstrictor that reduces local blood flow, directly into the hippocampus (Faraji et al., 2014; Fluri, Schuhmann, & Kleinschnitz, 2015; Sharkey, Ritchie, & Kelley, 1993; Sommer, 2017). These studies have shown that endothelin-1 leads to both subtle and overt cognitive decline as well as some cell loss, depending on timing of measures post-injury (Driscoll, Hong, Craig, Sutherland, & McDonald, 2008; Faraji, Metz, & Sutherland, 2011; Faraji et al., 2014; Farokhi-Sisakht et al., 2020; Mateffyova, Otahal, Tsenov, Mares, & Kubova, 2006; McDonald, Craig, & Hong, 2008; Sheng et al., 2015). As far as we are aware, only one study has demonstrated acute effects of hippocampal SI on cognitive decline (Sheng et al., 2015). There is a wealth of evidence as to the importance of gliosis, inflammatory pathway activation, and cell death in the pathophysiological consequences of ischemia (Brouns & De Deyn, 2009). However, it is unknown whether these cellular processes also underlie the cognitive outcomes of hippocampal SI. Thus far, the literature has relied on the hypothesis that the cellular processes underlying SI are similar to those observed in traditional ischemia models. However, in the absence of evidence, this hypothesis remains purely speculative. It is critical to establish whether similar processes underly the effects seen in hippocampal SI in order to begin to establish a clear model of the disorder along with potential treatment targets. Further, it is important to establish early consequences of hippocampal SI in order to identify clear targets for treatment.

To address this gap in the literature, the current study used a rat model to examine the immediate, short-term cellular processes and cognitive consequences of hippocampal SI. Rats received a unilateral infusion of endothelin-1 into the CA1 region of the dorsal hippocampus. Importantly, the use of unilateral lesions mirror the human condition, which is characterized by the presence of focal unilateral lesions (Leary & Saver, 2003). Twenty-four hours later, cognitive dysfunction was examined using the object place recognition task, a hippocampal-dependent spatial memory task (Olton, Becker, & Handelmann, 1979). To examine the cellular responses underlying the effects of hippocampal SI on cognition, we selected markers known to be common to both ischemic injury and dementias (Brouns & De Deyn, 2009; Deb, Sharma, & Hassan, 2010; Heinonen et al., 1995). We used cresyl violet staining to assess cell loss and immunohistochemistry to assess changes in synaptophysin, gliosis, apoptosis and inflammation in the hippocampus.

## 2. Materials and Methods

### 2.1. Animals

Twenty experimentally naïve, male Sprague-Dawley rats (380 - 410g) obtained from Animal Resources Centre, Perth, Western Australia. Rats were housed in plastic cages (67cm x 40cm x 22cm) located in colony room maintained on a reverse 12:12 light/dark cycle. There were four rats per cage with *ad libitum* access to chow and water. The experiment was approved by the Animal Care and Ethics Committee of UNSW Australia (ACEC 19/76A), performed in accordance with the Australian National Health and Medical Research Council’s ethical code, and reported in line with the Animal Research: Reporting In Vivo Experiments (ARRIVE) guidelines (Percie du Sert et al., 2020).

### 2.2. Induction of Hippocampal Silent Infarct

Rats were randomly allocated to either sham or SI groups and either right or left hemisphere stereotaxic surgery. Animals were anesthetized using isoflurane (1.5%) in oxygen and a cannula was stereotaxically lowered into the left or right dorsal hippocampus (4.5mm anterior, 3.0mm lateral to bregma and 2mm ventral) (Figure 1).

**Figure 1.**
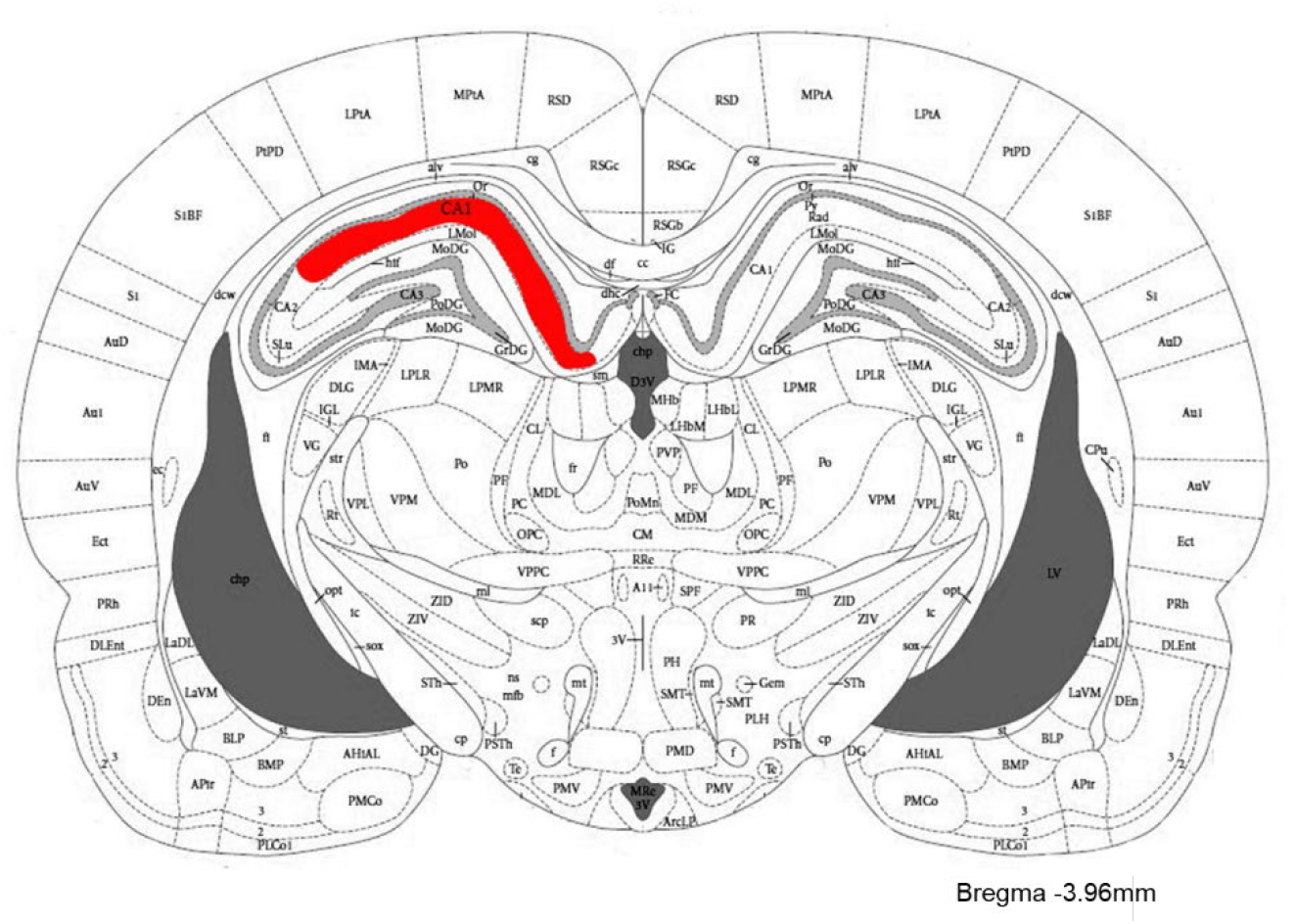
Representation from *The Rat Brain in Stereotaxic Coordinates* (Paxinos & Watson, 2006) atlas of the dorsal hippocampal CA1 targeted for unilateral stereotaxic surgery and subsequent cellular analyses. The left or right dorsal hippocampal CA1 as a target was counterbalanced between groups.

A 30-guage injector needle with a 1mm protrusion was inserted into the guide cannula and secured in place by a polyethylene tubing cuff. SI was induced using low dose of endothelin-1 (15pmol) dissolved in physiological saline (0.9% NaCl), for a total injection volume of 0.8μl. Previous research has shown that this dose of endothelin-1 reliably produces ischemic lesions in an adult rat model (Sheng et al., 2015). Rats in the sham group received a 0.8μl infusion of saline. Infusions occurred across two minutes and the injector was left in place for an additional five minutes prior to removal. After infusion, the injector and guide cannula were removed, the hole was covered with wax and the skin sutured. Rats were then returned to their home cage for 24 hours.

### 2.3. Object Place Recognition Task

To assess the consequences of hippocampal SI on memory, the object place recognition task was used as previously described (Finney et al., 2020). The task was performed in a square arena (60cm x 60cm x 50cm) made from wood and painted black with an oil-based paint. Prior to sham or SI surgery, rats were pre-exposed to the experimental arena twice, each for a period of 10 minutes, across two consecutive days. Following this pre-exposure, the object place recognition task was conducted to ascertain baseline performance. The task consisted of allowing the rats to explore two identical copies of commercially available objects (e.g. teacups) for a period of five minutes. They were then returned to their home cage for a five-minute retention interval. During this interval, the objects and arena were wiped with 80% ethanol and one object was moved to a new location on either the left or the right. The new location of the object was counterbalanced such that it was moved to the left for half of the rats in each group and to the right for the remainder. Following the retention interval, rats were returned to the arena and allowed to explore the objects during a three-minute test.

Subsequent to the baseline object place recognition task, rats underwent either sham or SI surgery. Twenty-four hours after surgery rats were again subjected to the object place recognition task as described above. The new location of the object was the opposite of the location to which the object had been moved at baseline (e.g. if object moved left at baseline it moved right at the 24-hour post-operative test) to ensure that rats did not use previous experience to perform the task.

A camera, mounted 1.85m above the arena was used to record rats’ exploratory behaviour during familiarization and test at both baseline and 24-hours post-operatively. Total time spent with both objects was calculated and exploration of the moved object was determined by the proportion of total time spent exploring the novel objects, such that: exploration proportion = (time_novel_ / time_novel_ + time_familiar_). Rats were considered to be exploring the object when at least their two back paws were on the floor of the arena and their nose was < 1cm from the object. Time with the objects was determined by manually scoring the video recordings using a MATLAB script (MATLAB, R2019b) with the scorer blind to the rat’s group allocation. Rats were excluded from statistical analyses if they spent < 0.5 seconds with one of the objects during either the familiarization or testing phases (Tran & Westbrook, 2015).

### 2.4. Hippocampal Tissue Collection

Immediately after the 24-hour post-operative behavioral test, rats were deeply anesthetized (150mg/kg pentobarbitone, intraperitoneal) and transcardially perfused with phosphate buffered saline (PBS) followed by 4% paraformaldehyde (PFA) in PBS. Whole brains were collected and post-fixed in 4% PFA for three hours followed by 30% sucrose for five days prior to sectioning. Forty μm thick free-floating sections were cut on a cryostat and collected in PBS at 4°C, then transferred to cryoprotectant solution (25% ethylene glycol, 25% glycerol in PBS) and stored at −20°C until use. Sections were collected from a 1 in 12 series (480μm apart, from 2.04mm to 5.52mm anterior to bregma).

### 2.5. Cresyl Violet Staining and Immunohistochemistry

To determine cell loss, four sections of the hippocampus (between 2.04 and 5.52mm anterior to bregma) were mounted onto microscope slides using 0.1% gelatin and stained with cresyl violet. Immunohistochemistry was also used to determine inflammation, synaptic loss and apoptosis in the hippocampus. Four hippocampal sections per rat were used. Sections were washed in PBS, incubated in 10mM, pH 6.0 sodium citrate antigen retrieval solution for 40 minutes at 70°C, and washed again in PBS. They were then blocked with a skim milk solution (5% skim milk powder, 0.1% bovine serum albumin, 2% serum, 0.3% Triton X-100 in PBS) for two hours at room temperature. Sections were then incubated for two nights at 4°C with primary antibodies against glial fibrillary acidic protein (GFAP; 1:5000; MAB360; Merck Millipore, USA), ionized calcium binding adaptor molecule 1 (Iba1; 1:1000; AB5076; Abcam, UK), synaptophysin (1:1000; AB14692; Abcam, UK); cleaved caspase-3 (1:1000; AB3623; Merck Millipore, USA), and interleukin 1β (IL1β; 1:200; AF-501-NA; R&D Systems, USA). All antibodies were made up in 1% serum, 0.1% bovine serum albumin, 0.3% Triton X-100 in PBS. Following incubation, sections were washed in static free solution (0.5% bovine serum albumin, 0.1% tween 20 in PBS) and PBS and a peroxidase block was used for 10 minutes (0.1% hydrogen peroxide in PBS). Sections were incubated in biotinylated secondary antibodies (1:1000; Vector Laboratories, USA) for two hours at room temperature. Secondary antibodies used included: horse anti-mouse (BA-2001), horse anti-rabbit (BA-1100), goat anti-rabbit (BA-1000), and rabbit anti-goat (BA-5000). Subsequently they were incubated with streptavidin horseradish peroxidase (HRP, 1:500 in PBS; Vector Laboratories, USA) for 2 hours. To visualize antigens, sections were incubated with 3,3-diaminobenzidine HRP substrate (Vector Laboratories, USA) for up to five minutes and mounted onto microscope slides using a gelatin solution (0.1% in PBS). Sections were then dehydrated in ethanol and xylene before being cover slipped using DPX mounting medium.

Stained slide-mounted hippocampal sections were imaged using a Vectra Polaris Automated Quantitative Pathology Imaging System (PerkinElmer, USA). Whole slide scanning was completed at 40x magnification and a resolution of 0.25μm / pixel. Images were viewed and analysed using QuPath v0.2.0 (Bankhead et al., 2017). One brain slice per rat was used for each antibody. The slice was taken from a region of the CA1 falling between −3.5mm and −4.5mm anterior to bregma (e.g. Figure 1) (Paxinos & Watson, 2006). Positively labelled cells in the CA1 region of the hippocampus were measured. Only the ipsilateral CA1 region of the hippocampus was included in analyses to control for damage from the stereotaxic cannulation surgery, independent of the endothelin-1 infusion. For cresyl violet and synaptophysin, the mean optical density of positive staining was recorded for each rat. Caspase-3 was quantified by counting the number of positively stained cells for each rat. For IL1β, GFAP and Iba1, a positive proportion of positively stained pixels was calculated such that: positive proportion = total number of positively stained pixels / total number of pixels. This ensured that expression of the respective protein along the cellular processes were adequately captured by the analysis. The mean positive proportion was recorded for each rat.

### 2.6. Statistical Analyses

All analyses were performed by an experimenter who was blinded to the experimental group. One rat from the SI group was excluded from all analyses due to death during surgery. Two additional rats from the SI group were excluded from the histological and immunohistochemical analyses due to sample damage during sectioning. Rats from either group were also excluded from histological and immunohistochemical analyses if there were inadequate samples, for example, due to problems with mounting or staining. If a rat failed to have an adequately stained sample from at least two of the six markers, it was excluded from all analyses. This resulted in one rat from the sham group being excluded. Data are represented as mean + standard error of the mean (SEM). Outliers were detected using the Grubb’s test, *p* < 0.01. To assess differences between the sham controls and hippocampal SI groups, independent samples t-tests were used followed by Cohen’s *d* to determine effect size. When multiple comparisons were made, the Holm-Sidak multiple t-test was used (Holm, 1979). To assess the relationship between time spent with the two objects on the object place recognition task, a simple linear regression was used. All statistical analyses were performed in GraphPad Prism (GraphPad Prism Software Inc., USA). Principal component analysis (PCA) was used to make inferences about the relationship between, and relative contribution of, the variables and their contribution to between-group differences. PCA is an unsupervised learning method that allows for interpretation of possible trends and patterns within an entire dataset (Lever, Krzywinski, & Altman, 2017). PCA was performed using R 3.6.3 (RStudio Inc., USA) package “factorextra”.

## 3. Results

### 3.1. SI leads to subtle cognitive impairment on the object place preference task

Prior to the hippocampal SI or sham surgery, rats underwent a baseline measure of their performance on the hippocampal-dependent object place recognition task. There was no significant difference between the groups at baseline, *t*(9) = 0.02521, *p* = 0.9804. At 24 hours after surgery, there were also no significant differences between groups with respect to exploration proportion during the three-minute test, *t*(9) = 1.434, *p* = 0.185. However, compared to their baseline exploration proportion prior to surgery, SI rats had a significantly decreased exploration proportion after injury, *t*(10) = 3.731, *p* = 0.008, *d* = 2.092 (Figure 2A). In contrast, there was no significant difference between baseline and post-operative performance on the three-minute test in the sham rats, *t*(8) = 0.8715, *p* = 0.409 (Figure 2A). To assess whether there were any locomotor differences between the groups on the 24-hour post-operative test, we examined the distance travelled by the rat during the three-minute testing phase. There were no significant differences between the groups, *t*(9) = 0.6746, *p* = 0.517 (Figure 2B).

**Figure 2.**
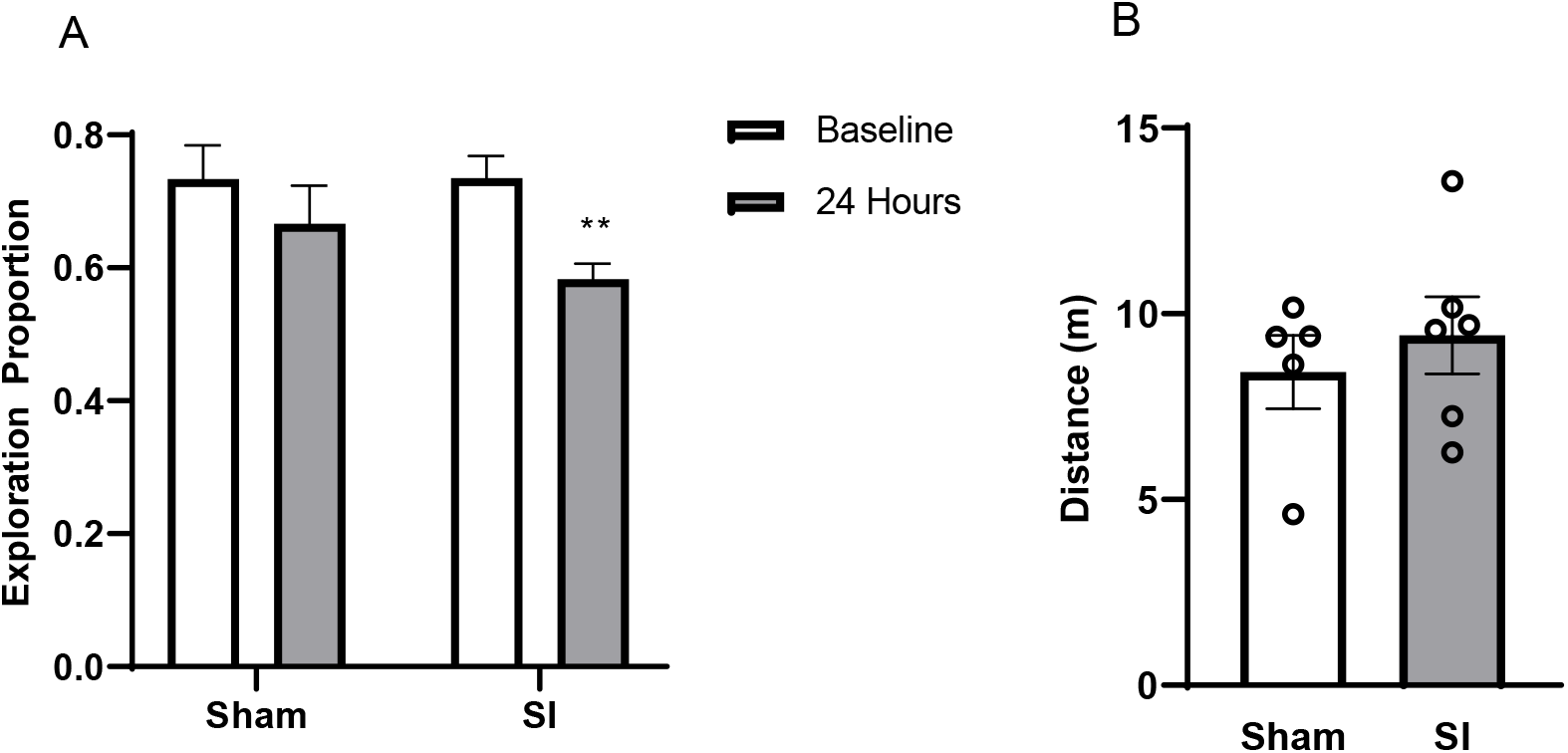
Performance on the object place preference task. (A) Exploration proportion during a three-minute testing phase at baseline and 24 hours after sham surgery or SI. Data expressed as mean + SEM. Data were analyzed using a Holm-Sidak multiple t-test. ** SI baseline vs. SI 24 hours, *p* < 0.01. (B) Distance travelled during a three-minute testing phase at 24 hours after sham surgery or SI. Data expressed as mean distance travelled (in meters) +SEM of sham and SI rats (n = 5-6 per group). Data analysed by independent samples t-test.

Further, 24 hours after SI, a simple linear regression analysis indicated that the SI rats performed the three-minute test differently than the sham group. Specifically, the more time that SI rats spent with the moved object the more time they spent with the unmoved object, *p* = 0.006 (Figure 3B), suggesting that they checked the objects continually during the task. This effect was not seen in the sham group, *p* = 0.630 (Figure 3A).

**Figure 3.**
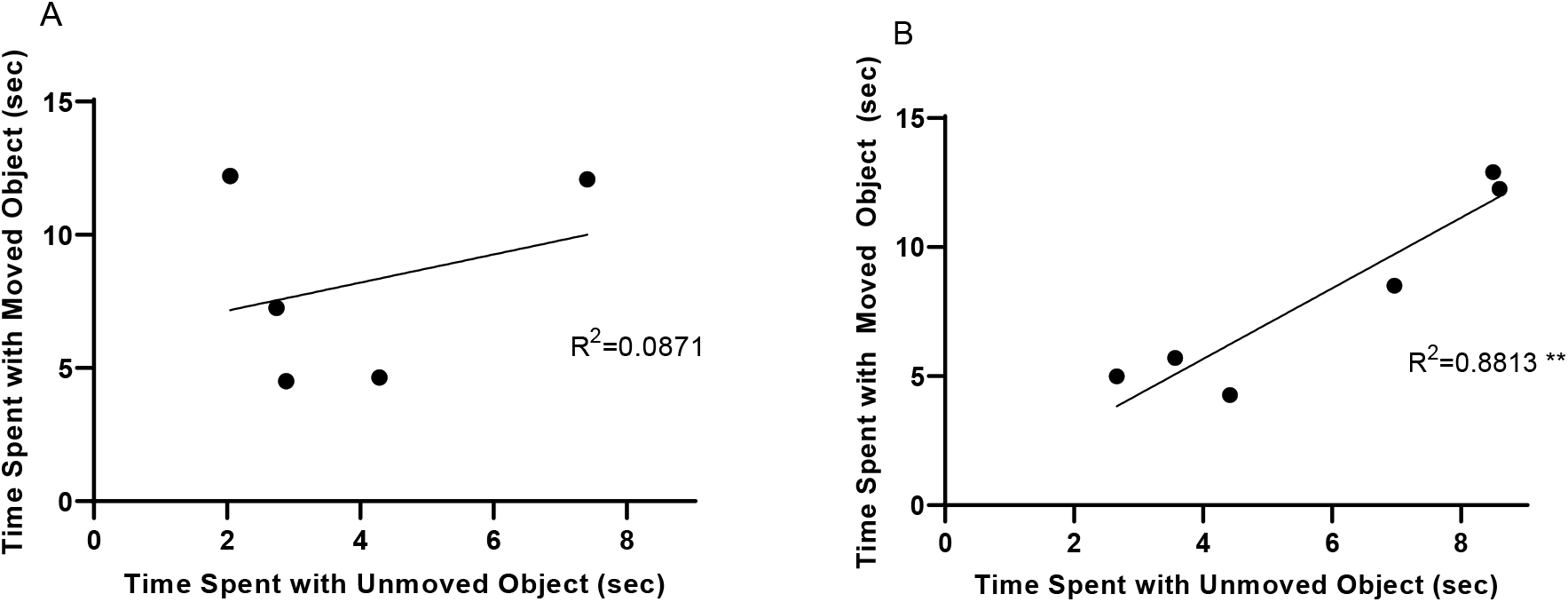
The relationship between time spent with the moved object and time spent with the unmoved object during a three-minute testing phase of the object place recognition task 24 hours after sham (A) or hippocampal SI (B). Simple linear regression analysis; ** *p* < 0.01.

### 3.2. SI does not lead to cell or synaptophysin loss but does increases apoptosis in the CA1 region of the dorsal hippocampus

Cresyl violet staining was used to determine whether hippocampal SI led to cell loss in the CA1 region of the dorsal hippocampus. The CA1 pyramidal layer appeared to be more densely packed in the sham than SI rats but there were no differences with respect to cell loss (Figure 4A). The statistical analysis failed to detect a significant between-group difference in cresyl violet optical density, *t*(9) = 1.461, *p* = 0.178 (Figure 4B). To assess pre-synaptic vesicles in the CA1, we used synaptophysin staining and compared optical density of the positively stained area between the groups. There appeared to be no differences in the pattern of synaptophysin staining throughout the CA1 of sham and SI rats (Figure 4A). This was confirmed by statistical analysis which failed to detect significant between-group differences in synaptophysin staining in the CA1 region of the dorsal hippocampus, *t*(9) = 0.660, *p* = 0.526 (Figure 4C). Despite a lack of cell and synaptophysin staining loss, we reasoned that cells may have been undergoing slower cell death processes beyond our 24 hour timepoint. Therefore, we assessed whether cells were undergoing apoptosis 24 hours after SI using immunohistochemical staining to determine the number of caspase-3 positive cells in the CA1. There were differences in the staining pattern of caspase-3 between the groups. Very few cells in the CA1 of sham rats were caspase-3 positive. In contrast, there were many positively stained caspase-3 cells in the CA1 of SI rats (Figure 4A). The statistical analysis confirmed that there were a significant increase in caspase-3 positive cells in CA1 of SI rats relative to sham controls, *t*(10) = 4.933, *p* = 0.0006, *d* = 2.848 (Figure 4D).

**Figure 4.**
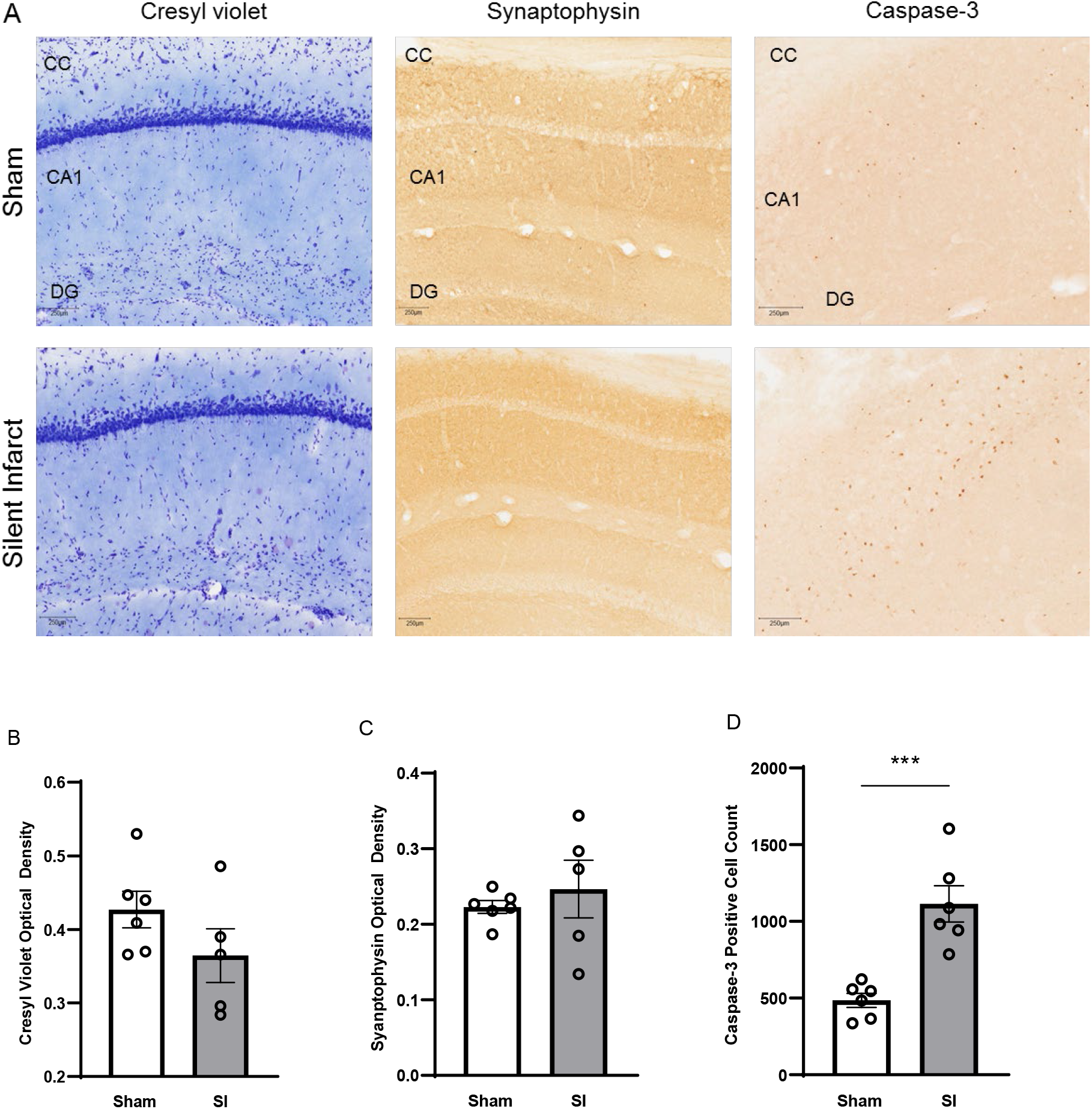
Hippocampal SI does not decrease the number of remaining cells or synaptophysin staining in the dorsal hippocampal CA1 region but does increase caspase-3 24 hours after injury. (A) Representative images of cresyl violet, synaptophysin and caspase-3 staining in the CA1 of sham and SI rats. Scale bar = 250μm. CC: corpus collosum; DG: dentate gyrus. (B) Cresyl violet and (C) synaptophysin in the CA1 were quantified using optical density. (D) Caspase-3 in the CA1 was quantified by counting the number of positively stained cells. All data expressed as mean + SEM of sham and SI rats (n = 5-6 per group). Independent samples t-test; *** *p* < 0.001.

### 3.3. SI increases gliosis and inflammation in the CA1 region of the dorsal hippocampus

To assess whether hippocampal SI led to increased gliosis in the CA1 dorsal hippocampal region, we stained for microglia, using the marker Iba1, and astrocytes, using GFAP. Both stains were quantified using a positive proportion measure to account for morphological changes characteristic of these cells following injury. There was a significant increase in Iba1 staining at 24 hours after injury in the hippocampal SI rats relative to the sham controls, *t*(10) = 2.518, *p* = 0.031, *d* = 1.492 (Figure 5B). As indicated by the staining pattern, this was reflected in increased number, density and size of the microglia present after SI relative to sham. Further, the sham rats appeared to have predominantly normal ramified microglia characterized by small cell bodies and longer processes, whereas in SI microglial cells appeared thicker and more densely stained, which is indicative of a reactive phenotype in response to injury (Figure 5A). There was also a significant increase in GFAP staining in the CA1 in SI relative to sham rats, *t*(9) = 3.003, *p* = 0.015, *d* = 1.915 (Figure 5C). The increase in GFAP staining following SI was dramatic, with increased number, size and density of astrocytes compared to sham. Additionally, glial reactivity in the sham rats appeared to be minimal whereas in the SI group there was indication of glial cell proliferation and hypertrophy (Figure 5A). To determine whether the increase in reactive gliosis was associated with an increased in glial pro-inflammatory cytokines, IL1β was used. As IL1β is commonly expressed by glial cells, the positive proportion measure was used to account for staining along microglial and astrocytic cellular processes. There was a significant increase in the amount of IL1β in the CA1 dorsal hippocampal region after hippocampal SI relative to sham controls, *t*(9) = 3.471, *p* = 0.007, *d* = 2.190 (Figure 5D). This increase in IL1β staining can be seen throughout the CA1 region of hippocampal SI relative to sham (Figure 5A).

**Figure 5.**
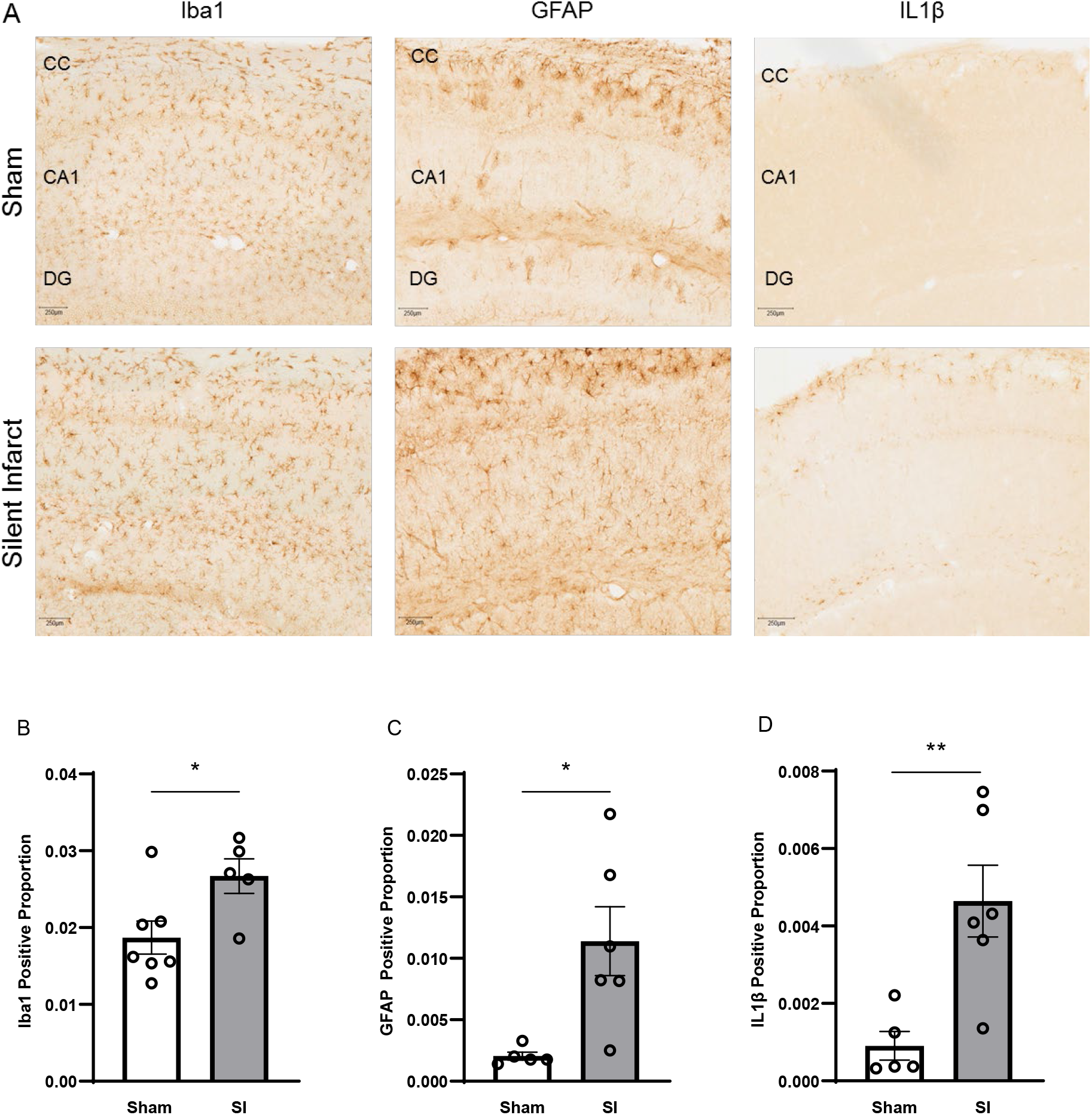
Hippocampal SI increased microglia (Iba1), astrocytes (GFAP) and the pro-inflammatory cytokine interleukin 1β (IL1β) in the dorsal hippocampal CA1 region 24 hours after injury. (A) Representative images of Iba1, GFAP and IL1β staining in the CA1 of sham and SI rats. Scale bar = 250μm. CC: corpus collosum; DG: dentate gyrus. (B) Iba1, (C) GFAP and (D) IL1β were quantified using positive proportion (positive pixels / total pixels) of positively stained pixels. All data are expressed as mean + SEM of sham and SI rats (n = 5-7 per group). Independent samples t-test; * *p* < 0.05, ** *p* < 0.01.

### 3.4. Principal component analysis suggests that gliosis and inflammation mediate the subtle cognitive decline seen in hippocampal SI

We used PCA to assess the relative contribution of and the relationship among the variables described above. There were nine variables of interest. Three of the measures were behavioural, based on performance during the object place recognition three-minute test 24 hours after injury: exploration proportion, time spent with the moved object and time spent with the unmoved object. The remaining six measures were based on histological and immunohistochemical staining, including: number of remaining cells (cresyl violet), pre-synaptic vesicles (synaptophysin), microglia (Iba1), astrocytes (GFAP), apoptosis (cleaved caspase-3), and pro-inflammatory cytokine (IL1β).

Two principal components accounted for 60.4% of the variance in the dataset. The PCA biplot showed that there was distinct clustering of the hippocampal SI and sham rats, with no overlap between them (Figure 6). The differences seen in the hippocampal SI rats were driven primarily by principal component 1 (horizonal axis, Figure 6): high levels of microglia, astrocytes, inflammation and apoptosis, as well as a greater amount of time spent with the unmoved object. Further, the SI rats were characterized by low levels of exploration proportion and lower number of cells. The sham rats, on the other hand, were characterized by high levels of exploration proportion, high cell numbers and correspondingly low levels of gliosis, inflammation, apoptosis and less time spent with the unmoved object. The time spent with the moved object during the object place recognition test and levels of pre-synaptic vesicle staining accounted for some of the variability in the dataset (principal component 2) but did contribute to the distinct clustering between the groups. Given that the PCA established and maintained clear clustering between the groups, we performed further statistical analyses on the significance of principal component 1 to the group distinction. A Welch two-sample t-test indicated that principal component 1 significantly contributed to the between-group differences, *t*(12.996) = 6.3013, *p* < 0.00001, *d* = 3.232.

**Figure 6.**
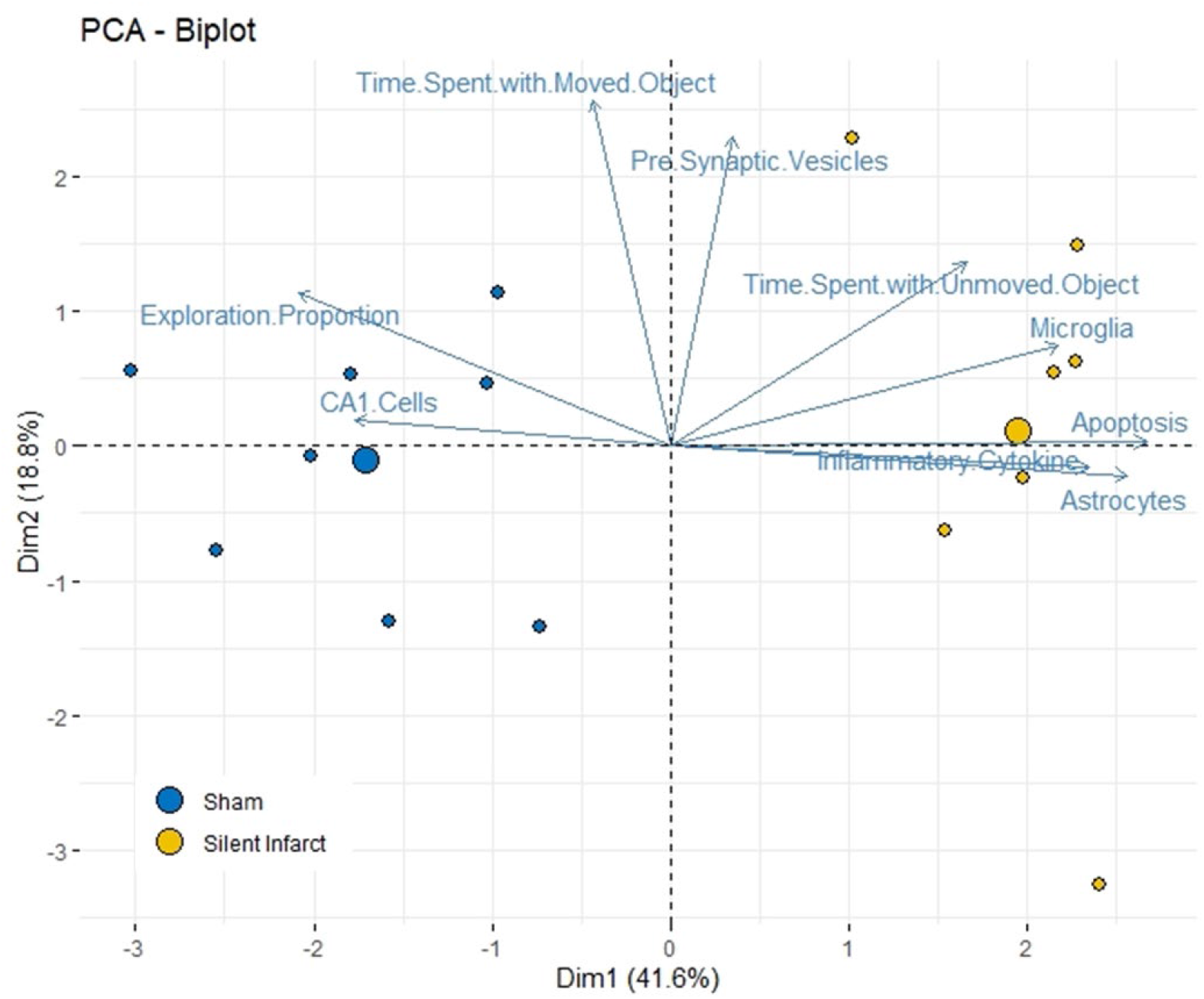
Principal component analysis biplot comparing principal components (Dim) 1 and 2.

## Discussion

The present study sought to determine the behavioral and pathophysiological consequences of hippocampal SI at 24 hours after injury in a rat endothelin-1 model. As far as we are aware, little is known about the short-term effects of hippocampal SI; for example, whether they are the same as those seen in other infarct models, such as ischemia. Thus far, the literature has relied on conjecture and speculation rather than evidence. Using the hippocampal-dependent object place recognition task, we found that there was a subtle cognitive decline in the SI rats relative to sham controls that was not related to any differences in locomotor behavior. This decline was characterized by significant differences in exploration proportion relative to baseline and a strong positive correlation between time spent with the moved and unmoved objects, suggesting that the SI rats continuously checked the two objects. Histopathological examination of the CA1 dorsal hippocampal region indicated that, while hippocampal SI did not lead to cell or synaptophysin staining loss, there was increased apoptosis, gliosis and inflammation. A PCA was used to identify trends in the dataset to establish relationships between, and relative contribution of, the various measures to hippocampal SI and sham groups. The PCA revealed distinct clustering of the SI and sham rats. Further, variance associated with hippocampal SI rats was primarily driven by higher levels of apoptosis, inflammation, gliosis and more time spent with the unmoved object during the object place recognition test. Importantly, spending more time with the unmoved object on the task indexes worse performance and suggests that the SI rats were impaired in differentiating between the object in the old location and the one moved to the new location, a phenomenon linked to worse memory and cognition. Taken together these results suggest that, as in the case of ischemia, early cellular responses of inflammation, gliosis and apoptosis in the dorsal hippocampal CA1 region after hippocampal SI underlies the subtle cognitive decline seen in these rats 24 hours after injury.

SI in people is associated with cognitive impairment and later development of dementias, including Alzheimer’s and vascular dementia (Blum et al., 2012; Vermeer et al., 2007). Hippocampal SI in rats is also associated with cognitive decline. Endothelin-1 infusions into the hippocampus resulted in deficits in contextual fear conditioning and the novel object recognition task 24 hours later (Sheng et al., 2015), however cell loss was not determined. Most rodent studies have examined long-term cognitive consequences, showing that SI leads to deficits in performance in the Morris Water Maze from five days up to three months after injury (Faraji et al., 2011; Farokhi-Sisakht et al., 2020; Li, Wang, Zhang, Fang, & Zhou, 2011; Mateffyova et al., 2006; McDonald et al., 2008; Mundugaru et al., 2018). Hippocampal SI decreased discrimination index on the novel object recognition task four weeks after stroke (Farokhi-Sisakht et al., 2020) and led to deficits on the Ziggurat task, a spatial memory task (Faraji et al., 2014). Here, we found that hippocampal SI is associated with subtle cognitive decline 24 hours after injury. While there were no significant differences between SI and sham groups on the object place recognition task, there was a significant decline relative to baseline in the SI group but not in the sham controls. Further, in SI rats, there was a strong positive correlation between time spent with the moved and unmoved objects during the test, suggesting that SI rats needed to check, or confirm, the object’s status. In other words, while they are still able to recognize the moved object, doing so depends on them continually checking the unmoved object as well, indicating that the SI rats required greater effort to perform the task relative to their sham counterparts. This idea is consistent with reports that in the early stage of Alzheimer’s disease, when cognitive impairment is mild, patients often rely on compensatory strategies to overcome their memory lapses, such as applying more effort, continued checking and asking questions (Buckley et al., 2015; Dixon, Hopp, Cohen, de Frias, & Backman, 2010; Ripich, Fritsch, Ziol, & Durand, 2000). In addition to indicating the cognitive decline associated with hippocampal SI is subtle, these several results suggest the need for more sensitive behavioral measures when assessing cognitive effects of SI (Faraji et al., 2014; McDonald et al., 2008). The present study demonstrates the importance of a baseline measure of cognitive function in addition to looking beyond the single measure of exploration proportion to better understand how the SI rats were solving the task.

Despite the well-documented cognitive effects of hippocampal SI, few studies have examined their neurobiological underpinnings, especially soon after injury. In the present study, cresyl violet staining showed that there was no significant cell loss in the CA1 region at 24 hours after injury. One previous study demonstrated tissue loss in the cortex 24 hours following intracerebral infusion of endothelin-1; however, it used a much higher concentration of endothelin-1 (160pmol) than in the present study (15pmol) (Soeandy et al., 2019). Cell loss following a subthreshold infusion of endothelin-1 (7.5pmol) was minimal when measured four days later in three-month-old rats but was extensive in older, 24-month-old rats (Driscoll et al., 2008). However, this study lacked a sham control group, raising the possibility that there had been a degree of cell loss in the younger rats, albeit to a lesser extent than the aged ones. Longer-term studies of cell loss after hippocampal SI have found extensive cell loss in the CA1 and dentate gyrus two-to four-weeks after injury (Faraji et al., 2011; Li et al., 2011; McDonald et al., 2008). It is possible that cells are still present in the CA1 despite being no longer functional at 24 hours after hippocampal SI, suggesting the use of other markers of neuronal dysfunction, such as Fluoro-Jade B (Schmued & Hopkins, 2000).

Synaptophysin plays a critical role in long-term synaptic plasticity in the CA1 (Janz et al., 1999). Loss of synaptophysin in the hippocampus has been associated with cognitive impairment in Alzheimer’s disease and may even be an early indicator of the disease (Heinonen et al., 1995; Sze et al., 1997). However, at 24 hours after injury, we found no effect of hippocampal SI on synaptophysin, suggesting that, at this time point, neurons have retained pre-synaptic structure. This suggestion is further evidenced by the finding that, while producing subtle memory deficits on the object place recognition test, hippocampal SI did not abolish recognition memory, implying remaining neuronal function in the CA1 at 24h after SI. Previous literature on synaptophysin immunoreactivity after ischemic injury has been mixed. One study found a reduction in CA1 synaptophysin two days after global ischemia in gerbils (Ishimaru, Casamenti, Ueda, Maruyama, & Pepeu, 2001), whereas another study found increased synaptophysin levels three to seven days after focal ischemia in the rat striatum prior to declining and ultimately disappearing by one month (Korematsu, Goto, Nagahiro, & Ushio, 1993). Regardless, it may be the case that 24 hours is too early to detect any changes in synaptophysin.

We used cleaved caspase-3 immunoreactivity to assess whether cells in the CA1 were undergoing apoptosis and found that hippocampal SI led to a significant increase in cleaved caspase-3 24 hours later, indicating that cells in the CA1 were undergoing apoptosis. Caspase-3 is known to play a critical role in ischemia-induced apoptosis (Brouns & De Deyn, 2009; Deb et al., 2010). Previous evidence has similarly found that an endothelin-1 infusion into the cortex of mice significantly increased caspase-3 levels 24 hours later (Soeandy et al., 2019). Longer term studies have demonstrated that caspase-3 remains upregulated four weeks after injury (Farokhi-Sisakht et al., 2020), and, in a severe model that involved overexpressing endothelin-1 and inducing a stroke, animals had increased caspase-3 levels in the hippocampus seven days after injury (Zhang, Yeung, McAlonan, Chung, & Chung, 2013). It is clear from our study that cells in the CA1 are undergoing apoptosis 24h after SI.

Gliosis, marked by increased reactive astrocytes and microglia, and an increase in inflammatory cytokines, play well-documented roles in inflammation after ischemic injury (Brouns & De Deyn, 2009; Deb et al., 2010). However, none of these phenomena have been studied at the 24h timeframe after hippocampal SI. In the present study, we found that at 24 hours after hippocampal SI there was already a significant increase in reactive astrocytes (GFAP) and microglia (Iba1). Increased expression of GFAP throughout the hippocampus has been found at three weeks after a unilateral infusion of endothelin-1 into the dentate gyrus (Li et al., 2011). Using spontaneously hypertensive rats, who show small white matter lesions characteristic of SI, another study found that both GFAP and Iba1 were increased at 35 weeks (Gao et al., 2019). Increased gliosis is associated with ischemic stroke and neurodegenerative disorders including Alzheimer’s disease, especially when reactive gliosis continues in the long-term (Burda & Sofroniew, 2014; Czlonkowska & Kurkowska-Jastrzebska, 2011; Pekny & Nilsson, 2005; Pekny, Wilhelmsson, & Pekna, 2014). This continuing process worsens apoptosis and increases subsequent cell loss. Thus, gliosis could be one of the factors underlying the early apoptosis seen in the present study. Further, astrocytes contribute to increased expression of β-amyloid precursor protein (APP) mRNA around the penumbra and core region of the infarct area after focal ischemia (Shi, Yang, Stubley, Day, & Simpkins, 2000) which may be linked to delayed cognitive decline that is sometimes observed following injury. Importantly, overaccumulation of APP is implicated in the development of cognitive dysfunction and Alzheimer’s disease (Yanker, 1996; Zhang, Thompson, Zhang, & Xu, 2011) and may play a role in the cognitive consequences of the increased gliosis after hippocampal SI seen in the present study.

Microglia changes have not yet been examined in SI. In ischemia, although consistently shown to be rapidly upregulated and activated, microglia play paradoxical roles, facilitating both recovery and degeneration (Gehrmann, Bonnekoh, Miyazawa, Hossman, & Kreutzberg, 1992; Patel, Ritzel, McCullough, & Liu, 2013). Similar to findings observed in focal ischemia models, we found that SI increased microglia. Glial cells are also known to release the pro-inflammatory cytokine interleukin-1 after ischemia (Brouns & De Deyn, 2009; Deb et al., 2010). Glial cells respond rapidly at the onset of ischemic injury and become the main source of brain inflammation (Czlonkowska & Kurkowska-Jastrzebska, 2011; Deb et al., 2010). In line with this, we found an increase in the expression of IL1β in the CA1 24 hours after hippocampal SI. This is similar to another study that found increased IL1β levels throughout the hippocampus around 3 weeks following dentate gyrus SI (Li et al., 2011) and at 24 hours after ischemic stroke (Deb et al., 2010). Similar findings have also been noted in spontaneously hypertensive rats with white matter lesions, an increase which the authors ascribed to decreased cerebral oxygen levels after SI (Gao et al., 2019).

It is worth noting that while both astrocytes and microglia are linked to neurodegeneration, subtypes of each are also involved in neuroprotective responses to injury (Becerra-Calixto & Cardona-Gomez, 2017). It may be the case in the present study that the hippocampal-SI induced increase in astrocytes and microglia are neuroprotective in nature, rather than neurodegenerative. However, the corresponding increase in IL1β suggests otherwise. Specifically, in ischemic stroke, increased IL1β in the acute injury phase has consistently been shown to lead to worse outcomes post-stroke (Sobowale et al., 2016). Further, previous research has shown that glia expressing the pro-inflammatory cytokine IL1β contribute to neurodegenerative rather than recovery processes (Patel et al., 2013; Varnum & Ikezu, 2012). SI also increased IL1β levels in the current study, suggesting that the glia may have promoted early degenerative processes. Future research would benefit from a phenotypic analysis of astrocyte and microglia after hippocampal SI along with examining the co-localization of glia and IL1β to better determine if glial activation early after SI promotes cell death at later time points.

The present study employed a PCA to establish the relative importance of the variables in defining the characteristics of hippocampal SI. We found that hippocampal SI rats formed a unique cluster relative to the sham controls, suggesting clear differences between the two groups. The main drivers differentiating the two groups were increased levels of apoptosis, gliosis and inflammation in the CA1 in addition to increased time spent with the unmoved object, an indicator of cognitive dysfunction. Hippocampal SI was characterized by lower levels of cells in the CA1 and decreased exploration proportion. Combined, this PCA suggests that exploration proportion is dependent on the number of cells in the CA1. We did not find a significant effect of hippocampal SI on either measure suggesting, perhaps not surprisingly, that 24 hours after injury rats are still in the early stages of CA1 injury and cellular dysfunction. On the other hand, increased time spent with the unmoved object, an indication that rats are not able to recognize novelty and are therefore cognitively dysfunctional, was associated strongly with gliosis, inflammation and apoptosis in the CA1. This suggests that in the early stages of injury after hippocampal SI, the cognitive effects are primarily due to early degenerative processes marked by inflammatory responses and the beginnings of cell death. In a study of spontaneously hypertensive rats with white matter lesions characteristic of SI, increased pro-inflammatory cytokine levels were associated with long-term cognitive impairment at 35 weeks on the Morris Water Maze task (Gao et al., 2019). Future research would benefit from continuing to examine the role of inflammation and gliosis in the early cognitive dysfunction seen after hippocampal SI. Further, gliosis and inflammatory pathways may represent a suitable target for ameliorating the cognitive effects seen after subcortical SI in the hippocampus. Future studies should examine the effectiveness of potential treatments targeting these processes using similar pre-clinical models of hippocampal SI.

## Acknowledgements

The authors thank the Biomedical Imaging Facility and the University of New South Wales, Australia, for the use of their Vectra Polaris scanner and to Iveta Slapetova and her team for helping to scan the microscope slides. The authors also thank Artur Shvetcov for his invaluable advice on principal component analyses.

